# Quantitative assessment of cell fate commitment in single-cell transcriptomics using scCS

**DOI:** 10.64898/2026.07.27.741070

**Authors:** Emil Kriukov, Ekaterina Ivleva, Petr Baranov

## Abstract

Cell fate trajectory inference is one of the key downstream methods in single-cell RNA-sequencing data analysis. Multiple existing tools allow studying such and help identify continuous cell fate dynamics, important genes along the trajectory, and reconstruct the transcriptional states a cell passes to its estimated final state. These methods have been essential to study various biological systems, yet cell fate by itself currently presents mostly qualitative analysis, and existing tools do not allow to directly quantify the fate-related parameters, including commitment, transition speed, fate affinity and entropy. Such quantifications may be performed through multiple parameters and result in better understanding and description of cell fates.

We present scCS (single-cell Commitment Scoring), a scverse-friendly Python framework for this problem. scCS introduces Discounted Future-Fate Propagation (DFFP), which models the source transition graph as a geometrically stopped random walk that can reach endpoint anchors or stop unresolved, with a user-defined finite expected graph horizon. For each cell, the resulting probabilities are separated into total fate reach, relative affinity among reached fates, entropy-based fate specificity and reach-supported resolved commitment, while Signed Ordering Flux independently quantifies local progression. scCS also provides endpoint-anchor, graph-coverage and horizon-sensitivity diagnostics, an instantaneous local-direction mode, standardized visualizations, gene-level analyses and replicate-aware comparisons across experimental conditions. Applications to pancreatic endocrinogenesis and neural crest-Schwann-cell differentiation illustrate the general framework. scCS converts an explicit biological fate hypothesis into auditable cell-, population- and replicate-level quantities without redefining the source dynamics.

## Introduction

Single-cell transcriptomics provides high-dimensional snapshots of heterogeneous populations but does not directly observe the path followed by an individual cell. Trajectory-inference methods estimate continuous orderings and branching structures from similarities among measured states. Slingshot[1] and Monocle 3[2] infer lineage geometry and pseudotime, whereas RNA-velocity methods estimate directional change from molecular kinetics and can generate a directed cell–cell transition graph[3,4]. Palantir models differentiation as a stochastic process and reports pseudotime, terminal-state probabilities and differentiation potential[5]; CellRank uses Markov-chain analysis to identify macrostates and calculate fate probabilities[6]. Together, these approaches describe continuous biological process organization, direction and likely outcomes.

Biologically, any given continuous transition is not always a single deterministic path. Cells can occupy overlapping intermediate states, pause, reverse, circulate locally or remain connected to outcomes not represented in a selected branch[7]. A directed transition graph therefore describes a distribution of possible next states rather than one inevitable future. In this setting, commitment has at least two components: whether the candidate fates are reachable at all and, conditional on reaching them, whether the future is concentrated on one fate.

Many experiments also begin with prior biological knowledge. Cell annotations, lineage evidence, spatial context or perturbation design may already justify a shared root and a finite set of candidate fates. The relevant question is then not “what topology and terminal states are present?” but “what does the measured dynamical graph imply for this supplied fate model?” A supervised analysis should keep this distinction visible: the annotations define the question, the transition graph supplies the evidence and the display geometry should not replace the measured dynamics.

scCS represents cell-fate commitment using several complementary quantities rather than a single score. It distinguishes how strongly a cell’s future is connected to the supplied fates, which fate is favored, how clearly that future is resolved and whether the cell is moving forward or backward along an independently defined progression coordinate. This separation allows weak fate support, ambiguous fate choice and retrograde motion to be identified as distinct biological behaviors.

The same framework is extended to comparisons between experimental conditions while preserving the biological replicate as the unit of inference. scCS aggregates cell-level measurements within donors, animals, organoids or cultures and performs replicate-aware statistical testing, avoiding pseudoreplication caused by treating individual cells as independent samples[8].

scCS is implemented as a scverse-compatible Python package operating on AnnData objects. Users provide root and fate annotations, continuous ordering and a directed transition matrix, typically derived from RNA velocity. The package returns cell-and population-level commitment measurements, model and graph diagnostics, standardized visualizations, gene-level analyses and replicate-aware condition comparisons. Pancreatic endocrinogenesis and Schwann-cell differentiation are used as illustrative applications of the general framework.

## Results

### scCS maps a supplied fate hypothesis onto directed cell-state dynamics

scCS separates the biological question from the dynamical evidence (**Fig. 1A**). The user supplies one root population, two or more candidate fates and a continuous progression coordinate (pseudotime, latent time, CytoTRACE2[9] score, gene expression, pathway NES, etc.), whereas the data supply a directed cell-cell transition matrix produced through RNA Velocity based trajectory inference methods. The root and fate annotations define the supervised question; the ordering orients progression and identifies late cells that can serve as endpoint anchors; neither input rewrites the source transition graph. The same core model is exposed through SingleScorer for one dataset, PairScorer for two conditions and MultiScorer for three or more conditions, with shared outputs spanning cell-level metrics, population summaries, visualization, condition inference and gene-level interpretation.

**Figure 1.**
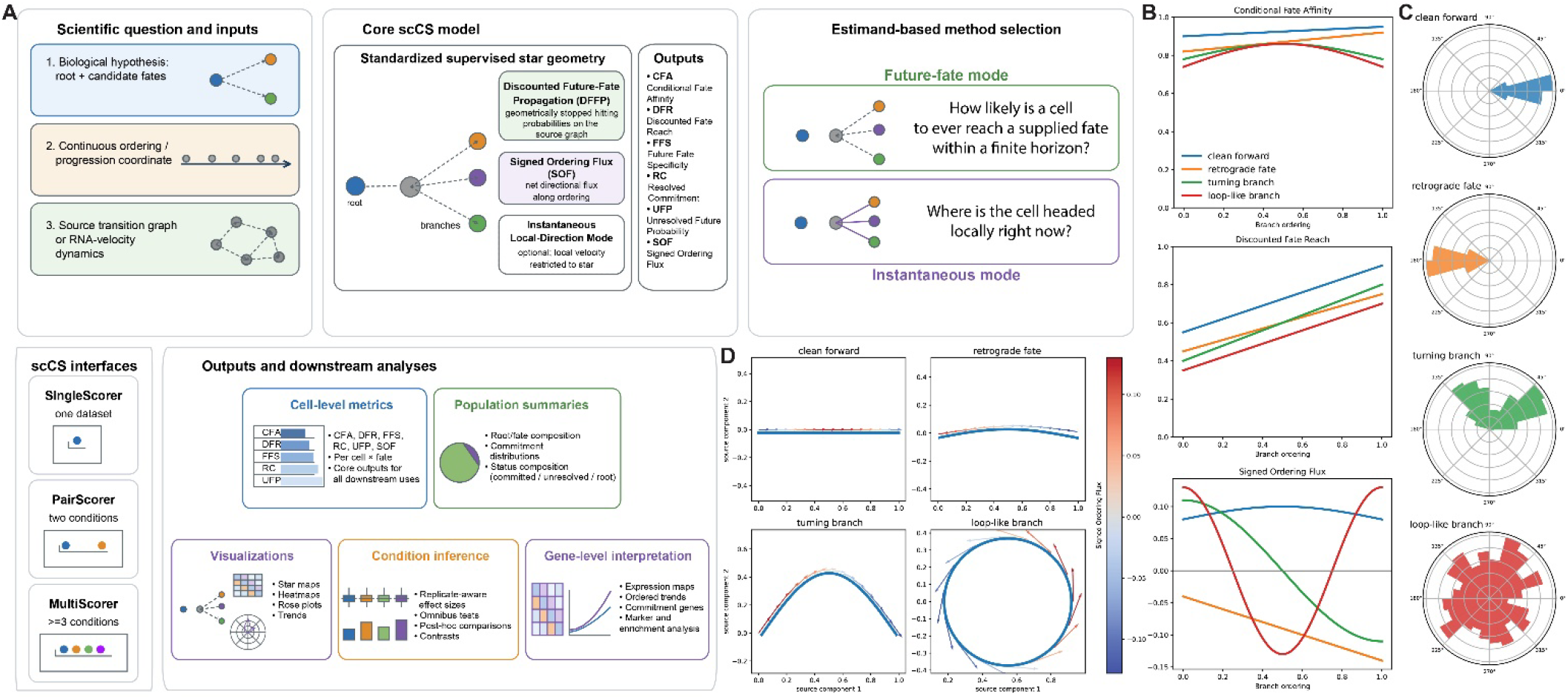
Overview of the scCS framework and separation of future-fate identity from local motion. a, scCS combines a supplied biological hypothesis (one root and candidate fates), a continuous progression coordinate and a directed source transition graph. The standardized star provides a common supervised geometry, while DFFP, Signed Ordering Flux (SOF) and the optional instantaneous local-direction mode return complementary cell-level quantities. SingleScorer, PairScorer and MultiScorer expose the same model for one, two or at least three conditions, respectively, and connect fitted scores to population, visualization, condition and gene-level analyses. The estimand is selected before interpretation: future-fate mode asks whether a cell can reach a supplied fate within a finite graph horizon, whereas instantaneous mode asks where the cell is directed locally. b, Synthetic clean-forward, retrograde, turning and loop-like branch archetypes shown as Conditional Fate Affinity, Discounted Fate Reach and SOF across branch ordering. c, Rose plots of the corresponding branch-relative velocity orientations. d, Source-space trajectories colored by SOF illustrate forward, retrograde, turning and loop-like motion. Panels b-d are controlled schematic examples rather than biological datasets.

Two estimands are kept distinct. Future-fate mode uses Discounted Future-Fate Propagation (DFFP) to ask how strongly a cell is connected to the supplied outcomes within a finite graph horizon, while preserving unresolved probability. Instantaneous mode asks where the cell’s one-step transition distribution is directed locally after it is represented in the supervised geometry (**Fig. 1A**). Thus, a long-range future association is not treated as equivalent to immediate motion.

Controlled branch archetypes illustrate this separation (**Fig. 1B-D**). Conditional fate affinity can remain high for a branch with clean forward motion, retrograde motion, turning or loop-like behavior, whereas Discounted Fate Reach increases with branch progression and Signed Ordering Flux distinguishes positive, negative and non-monotonic local movement (**Fig. 1B**). The corresponding rose plots summarize branch-relative velocity orientations (**Fig. 1C**), and source-space examples show that curved, reversing and loop-like trajectories cannot be reduced to one monotonic outward pattern without loss of information (**Fig. 1D**).

### scCS complements topology, velocity and fate-mapping methods

The closest computational alternatives solve related but different problems (**Table 1**). Slingshot[1] and Monocle 3[2] primarily reconstruct lineage geometry and pseudotime. scVelo estimates RNA velocity, kinetic parameters and latent time[4]; MultiVelo extends velocity modeling to paired chromatin and transcriptomic measurements[10]; RegVelo integrates gene-regulatory information with splicing dynamics[11]. Palantir[5] and CellRank[6] directly estimate probabilistic fates, with CellRank 2[12] supporting multiview transition kernels and macrostate discovery. scCS instead assumes that the root and candidate fate identities have already been justified and quantifies the consequences of that supplied hypothesis on a directed graph.

**Table 1.** Comparison of the primary estimands and built-in outputs of scCS and representative single-cell trajectory, velocity and fate-mapping methods. The table describes default or directly supported package outputs in the cited publications and documentation. “No” indicates that the quantity is not a native primary output of the cited workflow; it does not imply that a related analysis cannot be assembled by combining tools. RegVelo fate mapping is commonly performed by coupling its inferred dynamics to CellRank[6,11].

| Method | Primary task | Topology or terminal-state discovery | Per-cell probabilistic fates | Separates selected-fate reach from conditional identity | Explicit finite-horizon unresolved mass | Signed motion along a supplied ordering | Built-in biological-replicate condition inference |
| --- | --- | --- | --- | --- | --- | --- | --- |
| scCS | Supervised commitment quantification for supplied root and fates | No; supplied by user | Yes, for supplied outcomes | Yes | Yes | Yes | Yes |
| Palantir | Pseudotime, terminal fates and differentiation potential | Yes | Yes | No | No | No | No |
| CellRank 2 | Multiview Markov fate mapping and macrostate analysis | Yes or user-defined | Yes | No | No | No | No |
| scVelo | RNA-velocity, kinetics and latent time | Not primary | No | No | No | Indirect local direction | No |
| RegVelo | GRN-informed splicing dynamics and perturbation analysis | Not primary | Via downstream coupling | No | No | Indirect local direction | No |
| MultiVelo | Multi-omic chromatin–transcriptome velocity | Not primary | No | No | No | Indirect local direction | No |
| Slingshot | Lineage structure and lineage-specific pseudotime | Yes | Lineage weights, not future hitting probabilities | No | No | No | No |
| Monocle 3 | Principal-graph trajectory and pseudotime | Yes | No | No | No | No | No |

The comparison concerns built-in primary estimands rather than theoretical impossibility. Fate probabilities from another package can be analyzed externally, and user-defined terminal states can be supplied to other workflows. The distinctive combination in scCS is the explicit separation of total selected-fate reach from conditional identity, finite-horizon unresolved mass, signed progression along a separately supplied ordering and biological-replicate-aware condition inference within one package.

### Complementary metrics distinguish reach, identity, specificity and progression

For each cell, scCS stores fate-specific outcome probabilities and their derived metrics separately (**Table 2**). Discounted Fate Reach (DFR) is the total future probability assigned to the supplied fates. Conditional Fate Affinity (CFA) describes the relative distribution of that reached mass among the candidate fates. Future-Fate Specificity (FFS) measures how concentrated the conditional distribution is, Resolved Commitment (RC) combines reach with specificity, Unresolved Future Probability (UFP) retains mass not assigned to anchored outcomes at the declared horizon, and Signed Ordering Flux (SOF) reports one-step progression along the supplied ordering.

**Table 2.** Core scCS quantities and their interpretation.

| Quantity | Definition | Primary interpretation | Important caveat |
| --- | --- | --- | --- |
| **Outcome probability, pif** | Discounted probability assigned to selected fate f | Absolute support for a modeled future | Depends on graph, anchors and horizon |
| **CFA, qif** | pif divided by total selected-fate reach | Relative future identity among supplied fates | Conditional; unstable when reach is very low |
| **DFR, Ri** | Sum of selected-fate probabilities | How much future mass reaches any supplied fate | Not a measure of which fate is preferred |
| **FFS, Si** | One minus normalized entropy of CFA | Decisiveness among candidate fates | Can be high even when DFR is low |
| **RC, Ci** | DFR multiplied by FFS | Reach-supported specificity | Composite; report components |
| **UFP, Ui** | One minus all anchored outcome probabilities | Future not resolved at the declared horizon | Can include short horizon or omitted outcomes |
| **SOF, gi** | Expected one-step ordering change within selected cells | Forward, stationary or retrograde local progression | Depends on supplied ordering and path coverage |

This decomposition prevents distinct biological situations from being represented by the same scalar. High CFA with low DFR indicates that the resolved portion of the future favors one fate but the supplied outcomes have weak absolute support. High DFR with low FFS indicates that the outcomes are reachable but remain ambiguous. RC is high only when both reach and specificity are high, whereas SOF can be positive, near zero or negative independently of future-fate identity. Hard dominant-fate and status labels are convenience summaries and do not replace the continuous quantities.

### scCS integrates commitment scoring, local dynamics and gene-level readouts

We first applied SingleScorer to pancreatic endocrinogenesis[13], using endocrine progenitor states as the root and Alpha, Beta, Delta and Epsilon cells as candidate fates. The native RNA-velocity manifold provides the source dynamics (**Fig. 2A**), whereas the supervised star organizes the same selected cells around a shared root and symmetric fate directions (**Fig. 2B**). Transition-mediated vectors on the star retain the direction implied by the source transition operator rather than rebuilding neighbors in the display (**Fig. 2C**). Cell-level status labels distinguish fate-committed, future-ambiguous and low-reach states (**Fig. 2D**).

**Figure 2.**
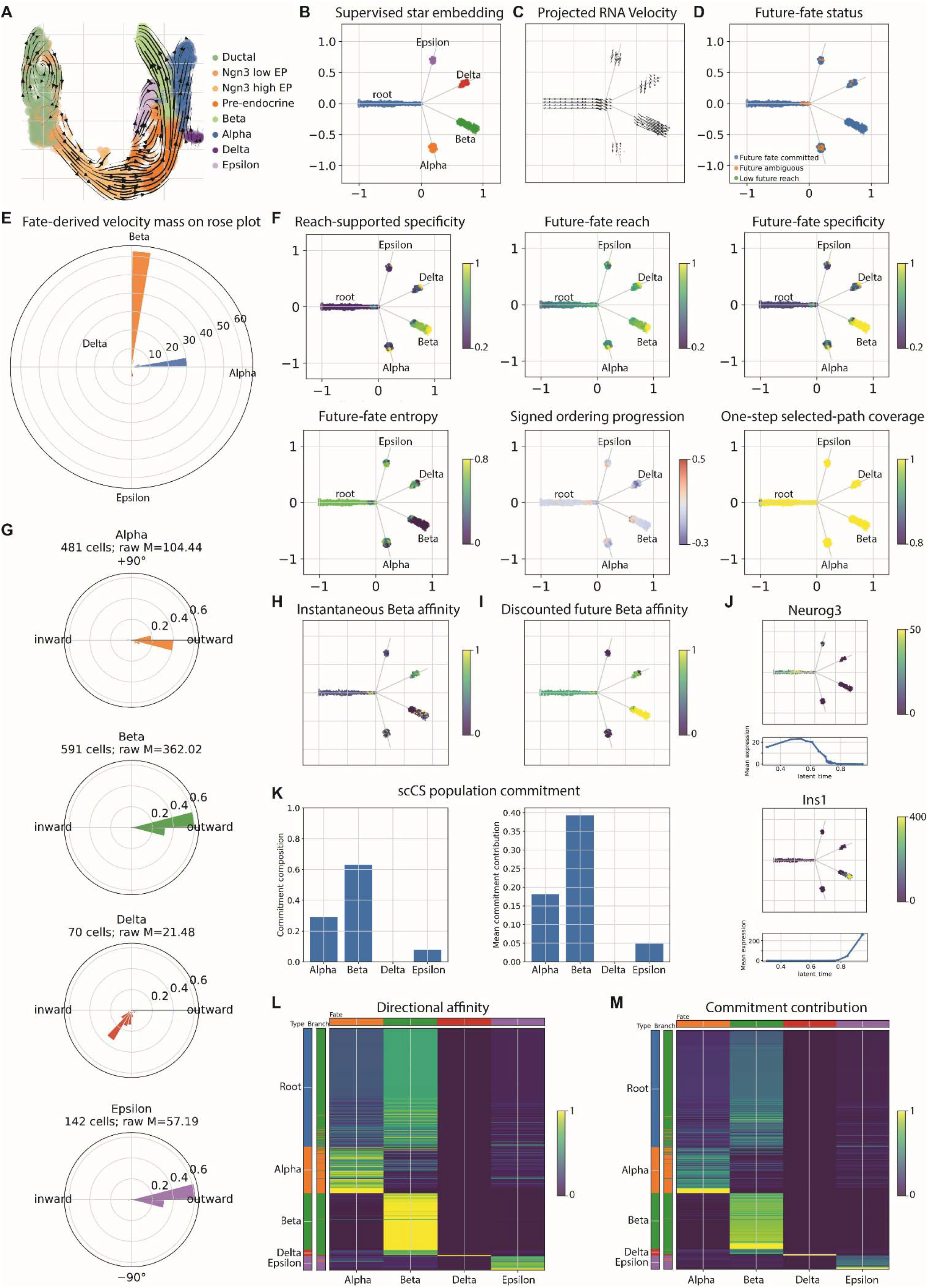
SingleScorer analysis of pancreatic endocrinogenesis. a, Native RNA-velocity manifold with annotated endocrine populations and streamlines. b, Standardized supervised star embedding of the selected root and Alpha, Beta, Delta and Epsilon fates. c, Transition-mediated RNA-velocity vectors recomputed on the star display. d, Cell-level future-fate status categories. e, Aggregate fate-derived velocity mass displayed as a rose plot. f, Star maps of reach-supported specificity, Discounted Fate Reach, Future-Fate Specificity, future-fate entropy, Signed Ordering Flux and one-step selected-path coverage. g, Branch-specific velocity-orientation profiles for Alpha, Beta, Delta and Epsilon; M denotes the raw fate-directed velocity mass reported by the plotting routine. h,i, Instantaneous and discounted future Beta affinity, respectively. j, Neurog3 and Ins1 expression on the star with mean expression trends along the supplied latent-time ordering. k, Population-level root-future composition and mean commitment contribution. l,m, Cell-level heatmaps of directional affinity and commitment contribution, with cells grouped by annotation and ordered within populations. The pancreas dataset is used as an illustrative application of the general SingleScorer workflow. Panels c, e, g and h show instantaneous transition-pushforward diagnostics, whereas panels d, f, i and k–m show DFFP-derived future-fate quantities. The two scoring modes were fitted to the same source transition graph but answer different questions.

The velocity summaries and fitted commitment metrics provide complementary views of the same furcation. The aggregate fate-directed velocity profile is dominated by the Beta direction, indicating that the largest share of instantaneous transition mass is aligned with the Beta branch in this dataset (**Fig. 2E**). The branch-specific rose plots refine this interpretation by showing that Beta and Alpha populations are characterized predominantly by outward-directed motion, whereas Delta and Epsilon exhibit weaker and less consistently outward profiles, including inward-oriented components (**Fig. 2G**). Thus, the annotated branches differ not only in the amount of velocity mass they receive, but also in whether local transitions are aligned with continued progression toward the corresponding fate.

The spatially resolved scCS metrics further separate several sources of heterogeneity across the furcation (**Fig. 2F**). DFR identifies regions whose future transition mass reaches the supplied terminal outcomes, whereas FFS and entropy distinguish cells with a concentrated fate distribution from those whose future remains divided among several alternatives. Reach-supported specificity combines these two properties, highlighting cells for which both fate access and fate resolution are strong. SOF adds an independent measure of local movement along the supplied ordering, revealing that cells with a relatively resolved future identity are not necessarily moving forward at the measured instant. Selected-path coverage provides an additional diagnostic by identifying regions where the local summary is supported by most of the outgoing transition mass and regions where a substantial part of the source dynamics lies outside the modeled furcation.

This distinction is particularly clear for the Beta lineage. Instantaneous Beta affinity highlights cells whose immediate transition vectors point toward the Beta direction, whereas discounted future Beta affinity identifies cells whose longer graph paths retain support for eventually reaching the Beta anchors (**Fig. 2H,I**). The two maps overlap in parts of the Beta branch but are not identical, especially among progenitor and non-Beta populations. These differences illustrate that a cell can be locally oriented away from Beta while still retaining discounted future-fate support at the declared effective horizon for a Beta future, or can transiently point toward Beta without having strong long-range reach to the Beta outcome. scCS therefore treats local direction, future reach and fate resolution as related but distinct components of commitment rather than combining them into a single trajectory score.

SingleScorer further connects the fitted commitment geometry to biologically interpretable gene-expression patterns and population-level fate structure. In the endocrine example, Neurog3 is enriched earlier along the supplied ordering and is concentrated in progenitor-associated regions of the star, consistent with an early endocrine differentiation program (**Fig. 2J**). By contrast, Ins1 rises later and is spatially restricted to the Beta branch, illustrating how mature fate-associated genes can be related directly to both branch identity and progression along the trajectory. These complementary views help distinguish genes associated with early state transitions from genes that mark later, fate-restricted differentiation.

Population-level summaries then condense the cell-wise probabilities into an interpretable description of the modeled root future (**Fig. 2K**). In this example, the root population is dominated by Beta-directed commitment, with smaller contributions toward Alpha and Epsilon and little contribution toward Delta. The corresponding mean fate contributions show that the Beta outcome is not only the most frequent conditional identity but also receives the largest absolute share of resolved future mass. This distinction is important because population composition and mean contribution summarize different aspects of the analysis: one describes how the modeled future is distributed among candidate fates, whereas the other reflects how strongly each fate contributes across cells.

The cell-level heatmaps retain this heterogeneity rather than reducing each population to a single average (**Fig. 2L,M**). Directional affinity shows a strong diagonal pattern, with annotated Alpha, Beta, Delta and Epsilon cells preferentially aligned with their corresponding fate directions. However, the off-diagonal signal indicates that some cells retain measurable support for alternative fates, particularly near transitional or incompletely resolved regions. Commitment contribution is more selective because it combines fate preference with the amount of future mass assigned to that fate. As a result, cells can display moderate affinity toward a fate but contribute little absolute commitment when their overall reach is low. Together, these heatmaps reveal both the expected population structure and the within-population variability that would be obscured by dominant-fate labels alone.

### MultiScorer allows multiconditional replicate-level comparison of cell fates

MultiScorer was evaluated on the neural crest-Schwann-cell dataset[11] using Common Progenitor as the root and Gut, Gut neuron and chromaffin-cell (ChC) populations as candidate fates (**Fig. 3A**). We used control, low and high pseudo-conditions to test condition-level recovery and treat these labels as software-validation scenarios rather than biological treatments. All conditions are represented on the same supervised geometry (**Fig. 3B**), so differences are interpreted under a common set of coordinates, anchors and fitted quantities.

**Figure 3.**
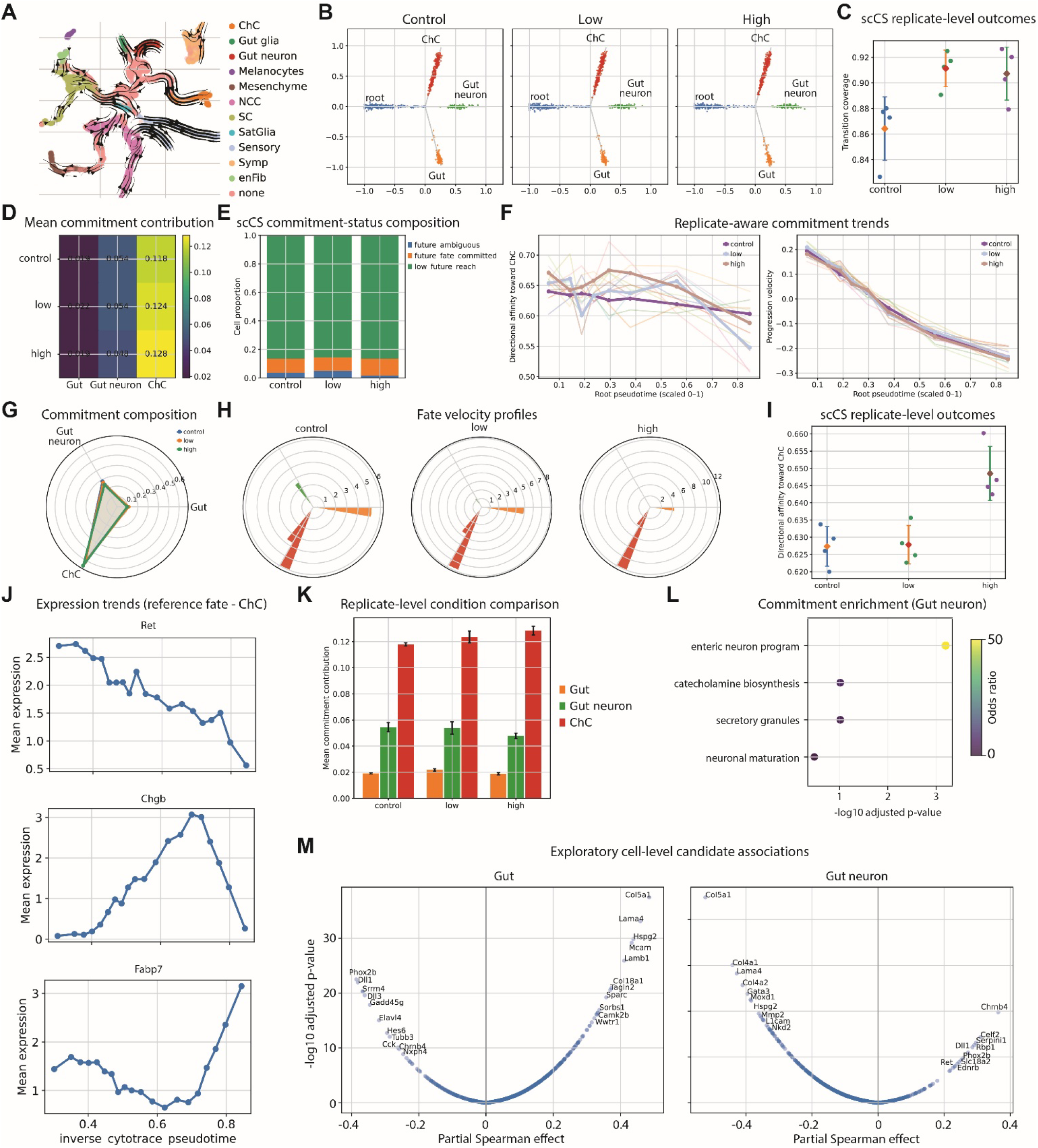
MultiScorer integrates shared geometry, replicate-aware condition comparison and downstream interpretation. a, Native dynamical RNA-velocity field for the neural crest-Schwann-cell dataset. b, Shared supervised star geometry for control, low and high pseudo-conditions. c, Replicate-level selected-transition coverage; points denote biological-replicate summaries and central markers with intervals summarize conditions. d, Mean commitment contribution to Gut, Gut neuron and ChC across pseudo-conditions. e, Composition of future-ambiguous, future-fate-committed and low-reach status categories. f, Replicate-aware trends in directional affinity toward ChC and progression velocity along scaled root pseudotime; faint lines represent replicate-level trends and emphasized lines represent condition summaries. g, Root-future commitment composition by condition. h, Condition-specific fate-velocity profiles. i, Replicate-level directional affinity toward ChC. j, Mean expression trends for Ret, Chgb and Fabp7 along inverse CytoTRACE2 score. k, Replicate-level comparison of mean commitment contribution. Bars denote means of the four pseudo-replicate summaries per condition, and error bars denote the standard error across pseudo-replicates. l, Over-representation analysis of Gut-neuron commitment-associated genes; point color denotes odds ratio and the horizontal axis shows -log10 adjusted P value. m, Exploratory partial Spearman associations between expression and Gut or Gut-neuron commitment after adjustment for the supplied ordering. Control, low and high are constructed tutorial labels used to validate the MultiScorer workflow and are not biological treatment groups. Pseudo-condition construction, replicate numbers, permutation settings and interval calculations are described in Methods. In panels c and i points represent constructed pseudo-replicate means, diamonds denote condition means and intervals show the condition mean ±1.96 standard errors calculated across the four pseudo-replicates.

Transition coverage remained consistently high across the constructed pseudo-replicates, indicating that most outgoing transition mass was retained within the modeled furcation and that condition-level differences were not driven by a loss of graph support in one pseudo-condition (**Fig. 3C**). Against this stable background, the fitted commitment summaries recovered the imposed condition structure. Mean fate contributions showed a progressive increase in ChC-directed commitment from control to the high pseudo-condition, accompanied by higher ChC directional affinity at the replicate level (**Fig. 3D,I,K**). Gut-neuron contribution changed more modestly and was slightly reduced in the high condition, whereas Gut contribution remained comparatively small. These results illustrate that MultiScorer can resolve fate-specific shifts even when the overall topology and transition coverage of the pooled model remain similar across groups.

The status-composition analysis provides a complementary population-level view (**Fig. 3E**). Most cells were classified as low reach in all three pseudo-conditions, with only a smaller fraction assigned as fate committed or ambiguous. The similarity of these proportions across conditions shows that an increase in commitment toward one particular fate does not necessarily require a large change in the global fraction of committed cells. Instead, condition effects can arise through redistribution of future support among candidate fates within an otherwise similar overall state composition. Reporting both fate-specific contributions and global status categories therefore prevents a localized lineage shift from being misinterpreted as a system-wide increase in commitment.

Replicate-aware trends further separated changes in future identity from changes in local progression (**Fig. 3F**). ChC directional affinity was higher in the high pseudo-condition across much of the root ordering, consistent with the fate-contribution summaries. By contrast, progression velocity declined along the ordering in all conditions with broadly similar trajectories. Thus, the imposed condition effect primarily altered the distribution of future support toward ChC rather than producing a general acceleration of movement through the root trajectory. This distinction illustrates why affinity and progression should be examined separately: cells may become more strongly associated with one fate without moving more rapidly toward it.

The same fitted framework supports condition-level visual and molecular interpretation. Root-future composition summarizes how the balance of modeled outcomes changes between conditions, while fate-directed velocity profiles show whether these changes are accompanied by altered local orientation toward Gut, Gut neuron or ChC (**Fig. 3G,H**). In this example, the ChC direction carries the strongest local velocity mass, whereas the Gut and Gut-neuron directions remain weaker. Comparing these profiles across control, low and high conditions helps distinguish a genuine redistribution of fate-directed motion from a change visible only in aggregated commitment probabilities.

Gene-expression trends provide an additional biological layer without changing the fitted commitment model. Ret, Chgb and Fabp7 follow distinct trajectories along inverse CytoTRACE2 score (**Fig. 3J**). Ret is highest earlier and declines with progression, Chgb increases toward intermediate-to-late states before decreasing, and Fabp7 rises more strongly at the later end of the ordering. These contrasting profiles illustrate how genes associated with early progenitor identity, neuroendocrine differentiation or later glial-like states can be positioned relative to the same continuous progression coordinate.

Downstream enrichment and gene-association analyses then connect variation in fitted commitment to candidate molecular programs. Genes associated with Gut-neuron commitment were enriched for an enteric-neuron program, together with neuronal maturation, catecholamine-biosynthesis and secretory-granule terms (**Fig. 3L**). Partial rank associations identified genes positively or negatively correlated with Gut or Gut-neuron commitment after adjustment for the supplied ordering (**Fig. 3M**). This adjustment is important because it separates association with commitment from simple co-variation along pseudotime. However, these analyses are performed at the exploratory cell level in this example. They should therefore be interpreted as illustrations of package functionality rather than as replicate-level tests of causal lineage regulators.

### Method evaluation defines the DFFP estimand, display limitations and computational regime

The method-selection experiments were organized around the biological and mathematical requirements of the target estimand rather than around visual preference. The synthetic source graph illustrates why this distinction matters: a biologically valid route from the root to a terminal anchor can leave the cells included in the displayed furcation, pass through an external state and later return before reaching the modeled outcome (**Fig. 4A**). A method that truncates the graph at the display boundary would treat this path as lost, even though it remains supported by the original transition structure. The formulation comparison therefore evaluates whether each approach preserves the source graph, allows leave-and-return paths, retains unresolved probability, accommodates curved or non-monotonic motion and exposes its assumptions through explicit parameters (**Fig. 4B**). Within this defined comparison, DFFP is the only formulation that satisfies all five requirements simultaneously. This result should not be interpreted as a universal ranking of trajectory or fate-mapping methods; rather, it identifies the formulation that is consistent with the specific supervised future-fate question posed by scCS.

**Figure 4.**
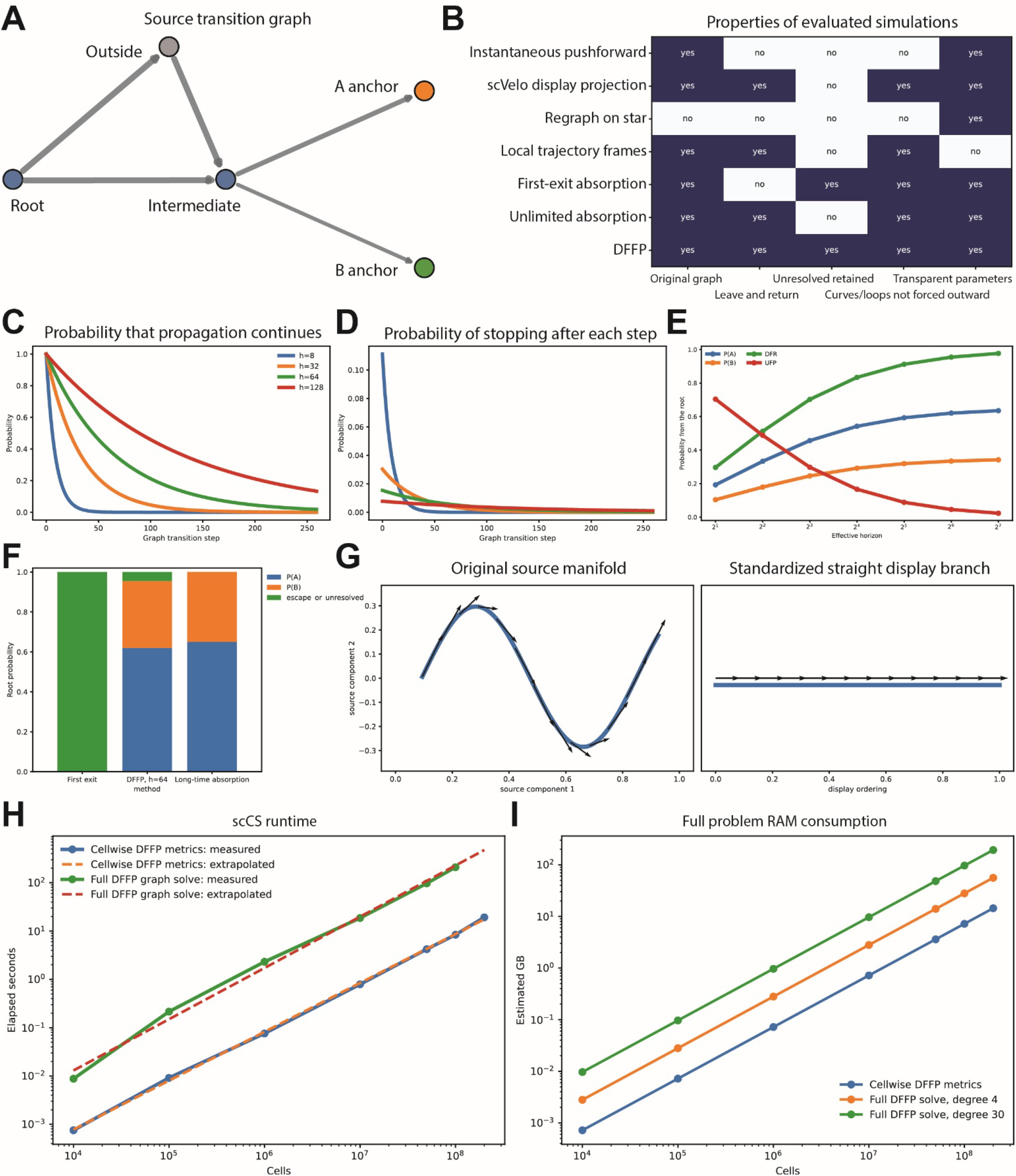
Methodological properties, sensitivity and computational scaling of DFFP. a, Synthetic source transition graph in which a valid root-to-anchor path leaves the selected furcation through an outside state and later returns. b, Property matrix for evaluated instantaneous, display-projection, regraphing, trajectory-frame and graph-propagation formulations. Columns indicate preservation of the original graph, support for leave-and-return paths, explicit unresolved mass, accommodation of curved or loop-like motion and transparent parameters. c,d, Probability that propagation continues and probability of stopping after each graph transition for effective horizons h = 8, 32, 64 and 128. e, Root probabilities for two anchors, total Discounted Fate Reach and Unresolved Future Probability across increasing effective horizons. f, Probability decompositions produced by first-exit absorption, DFFP at h = 64 and long-time absorption. g, A curved source manifold and its straight standardized display illustrate that display straightening cannot preserve every local source-space vector angle. h, Measured and extrapolated runtime for the cellwise DFFP metric transform and full DFFP graph solve. i, Analytic full-problem memory estimates for cellwise metrics and full DFFP solves at graph degrees 4 and 30. Panels a-g use controlled synthetic examples; panels h,i report the benchmark regimes described in Methods.

The effective horizon provides an explicit control over how deeply the random walk explores the transition graph. For small (h), propagation terminates rapidly, so the analysis emphasizes outcomes that can be reached within only a few graph transitions. As (h) increases, the probability of continuing remains substantial over a larger number of steps, whereas the probability of stopping at any particular step becomes smaller and more broadly distributed (**Fig. 4C,D**). The horizon therefore controls graph depth and outcome resolution, not developmental time in hours or days. It specifies how much indirect transition structure is allowed to contribute to the estimated future.

The effect of this parameter is visible in the synthetic fate probabilities (**Fig. 4E)**. At short horizons, a large fraction of the root future remains unresolved because the process often stops before reaching either anchor. Increasing (h) progressively transfers probability from UFP to the modeled outcomes, causing both fate-specific probabilities and total DFR to rise. The values approach a stable regime at longer horizons, but the unresolved component remains explicitly represented until the graph has been explored deeply enough to support an anchored outcome. This behavior makes horizon sensitivity biologically interpretable: instability across plausible horizons indicates that the fate assignment depends strongly on distant graph structure, whereas early stabilization suggests that the outcome is already supported by relatively local paths.

The comparison with first-exit and long-time absorption further shows that the three formulations produce different probability decompositions because they define different events (**Fig. 4F**). Under first-exit analysis, leaving the selected subset is classified immediately as escape, so the valid leave-and-return route in **Fig. 4A** cannot contribute to either fate. DFFP instead propagates on the complete graph and retains probability that remains unresolved at the declared finite horizon. Long-time absorption removes this finite-horizon uncertainty and assigns all mass that can eventually reach an anchor to the terminal outcomes. The differences among the bars are therefore expected consequences of the estimands, not numerical disagreement between equivalent methods.

The geometry experiment addresses a separate but related issue regarding the distinction between transferring dynamics and transforming an embedding. In the source manifold, a curved trajectory contains local vectors whose orientations change continuously along the path (**Fig. 4G**). Straightening the same trajectory into a standardized branch necessarily removes this curvature, and no single Euclidean rotation can preserve all original local angles. scCS therefore does not attempt to rotate UMAP or manifold arrows directly into the star. Instead, it applies the source transition operator to cell positions in the supervised geometry and recomputes the expected displacement there. The resulting arrows describe how the original transition relationships are expressed relative to the supplied root-and-fate hypothesis, while the native manifold remains the appropriate space for judging whether the underlying velocity field is credible.

Finally, the computational benchmarks separate two stages with distinct costs. Benchmarks evaluated simulated sparse transition-graph problems ranging from 10,000 to 200,000,000 cells. Cellwise metric calculation and the complete DFFP graph solve were benchmarked separately. Once DFFP outcome probabilities have been obtained, calculation of cell-level quantities such as CFA, DFR, FFS, RC and UFP is comparatively inexpensive and scales approximately linearly with cell number in the measured and extrapolated ranges (**Fig. 4H**). The complete sparse graph solve follows a similar overall scaling trend but has a substantially larger constant cost because it must operate on the transition matrix and maintain state-by-outcome solver arrays. The memory analysis shows the same distinction (**Fig. 4I**): cellwise metric calculation requires relatively little memory, whereas the full DFFP solve depends strongly on both the number of cells and graph degree. A degree-30 graph therefore reaches the host-memory limit much earlier than a degree-4 graph. Measured and extrapolated regimes are shown separately so that projected performance at very large cell numbers is not presented as directly observed computation.

## Discussion

scCS addresses a specific inferential gap between trajectory reconstruction and biological interpretation. It does not discover a trajectory, infer RNA velocity or determine terminal identities without supervision. Instead, it asks what a supplied root-and-fate model implies when evaluated against a directed transition graph and an independently supplied progression coordinate. This division of labor makes scCS naturally composable with scVelo[4], RegVelo[11], MultiVelo[10] or other methods that generate transition information, and with Slingshot[1], Monocle 3[2], Palantir[5] or CellRank[6,12] when their inferred structures are subsequently curated into a supervised biological hypothesis.

The central methodological choice is to preserve several quantities that are often collapsed. Conditional identity is not total resolution; specificity is not reach; and long-term future association is not local progression. DFFP introduces an explicit finite graph horizon and unresolved probability, while SOF measures signed movement along the supplied ordering (**Figs. 1B-D** and **4A-F**). Together, these outputs allow turning, loops, temporary reversals and omitted outcomes to appear in the analysis instead of being absorbed into a visually clean but biologically overdetermined branch.

### Projecting source dynamics into a supervised coordinate system

The transition-mediated projection in scCS separates the evidence describing cellular dynamics from the geometry used to interpret those dynamics. The source transition graph retains the directed relationships inferred in the native manifold, whereas the supervised geometry encodes the biological question specified by the user. scCS therefore transfers transition-weighted relationships between cells rather than attempting to rotate or warp arrows from UMAP or another nonlinear embedding. Such a direct geometric transformation would generally be non-unique because nonlinear embeddings do not provide an invertible map with a well-defined Jacobian between the source and supervised coordinate systems (**Figs. 1D** and **4G**).

Applying the same transition operator to the supervised coordinates asks where each cell’s outgoing transition distribution points relative to a declared root and set of candidate fates. This preserves the relational content of the dynamical model while deliberately replacing the metric structure of the original visualization. The regular-simplex construction gives every candidate fate an equal-length direction and places the root axis orthogonally to the fate subspace, preventing branch area, abundance or orientation in an unsupervised embedding from conferring an artificial geometric advantage.

The projected field nevertheless inherits the assumptions and errors of the source graph. Incorrect velocity estimates, batch-driven transitions or poorly connected neighborhoods remain incorrect when expressed on the star. Conditioning on the selected furcation can also be misleading when only a small fraction of outgoing transition mass is retained. scCS therefore reports transition coverage, suppresses vectors below a declared coverage threshold and treats the native manifold as the primary location for evaluating the credibility of the dynamical model.

The supervised star is intentionally a reduced scientific coordinate system. It does not preserve within-fate curvature or all local manifold structure, and projected arrows, rose plots and branch profiles should be interpreted as diagnostics of the source transition field under the supplied fate hypothesis rather than independent evidence that the hypothesis is correct. The projection is also distinct from DFFP: projected vectors describe immediate transition-induced motion, whereas DFFP propagates probabilities on the original graph to quantify finite-horizon future outcomes. Agreement between the two views can strengthen interpretation, while disagreement can reveal transient reversals, loops or cells that are locally directed toward one state but retain future support for another fate.

### Potential applications across development, disease and therapeutic intervention

scCS is applicable whenever a biologically justified root, a finite set of candidate outcomes, a progression coordinate and a directed transition model can be specified. In developmental and regenerative systems, it could quantify when progenitors acquire substantial reach toward particular lineages, whether fate preference precedes strong fate specificity and whether local progression agrees with longer-range future support. These questions complement lineage-tracing and Markov fate-mapping studies showing that transcriptional state can predict future outcomes without determining them uniquely[6,14].

Disease-associated state transitions provide another potential application. Microglia, for example, can move between homeostatic, inflammatory and disease-associated programs in neurodegenerative settings[15,16]. Our single-cell study of retinal myeloid cells after transplantation identified both homeostatic-to-activated and activated-to-homeostatic trajectories, suggesting that transplantation-associated microglial activation is at least partly reversible[17]. scCS could quantify whether an intervention reduces future reach toward maladaptive activation, increases support for recovery-associated states or changes local progression without changing longer-range state identity.

PairScorer and MultiScorer may also be useful for perturbation, drug-response and clinical cohort studies. Single-cell perturbation experiments and patient profiling increasingly reveal heterogeneous responses that cannot be represented adequately by mean expression changes alone[18,19]. scCS is capable of testing whether treatment redirects cells among prespecified outcomes, alters commitment strength or changes progression dynamics. Such analyses would remain replicate-level comparisons of a fitted state model and would require independent validation before being interpreted as predictors of patient benefit.

### Fate as an operational outcome rather than an assumption of irreversibility

In scCS, a supplied fate need not represent a permanently terminal lineage. In development, anchors may correspond to mature differentiated populations, whereas in inflammation, injury, cancer or treatment response they may represent reversible or metastable states. DFFP therefore measures discounted future-fate support at the declared effective horizon for reaching a defined outcome under the supplied transition model; it does not assert that the state is biologically irreversible.

This distinction is important for continuous states where velocity vectors may be connected in both directions. In these settings, terms such as *future-state support* or *state commitment* may be preferable to irreversible lineage commitment. Large UFP may indicate that the selected outcomes incompletely describe the response, that the horizon is too short or that cells remain connected to unmodeled intermediate states.

### Relationship to lineage tracing and experimental validation

scCS infers future-state support from molecular transitions and does not directly observe ancestry or descendants. Lineage tracing is therefore a complementary validation strategy. Combined lineage tracing and single-cell transcriptomics have shown that molecular state can be predictive of fate while remaining insufficient to determine it completely[14].

Where clonal or longitudinal data are available, anchors could be defined independently and scCS predictions compared with observed descendant frequencies. Time-course sampling, transplantation assays and genetic or pharmacological perturbations could similarly test whether predicted fate support changes in the expected direction. Gene associations and enrichment results generated by scCS should consequently be treated as candidate-generating evidence until the proposed regulators or transitions are validated experimentally.

### Core package limitations

scCS is a supervised framework, and its conclusions are only as credible as the biological model and dynamical inputs supplied to it. Root and fate annotations must therefore be well justified and sufficiently sampled, the progression coordinate must have a clear orientation and biological interpretation, and the transition graph should be inspected in the native manifold before commitment scores are interpreted. Endpoint anchors should also be evaluated rather than assumed to be ideal terminal states, because poorly supported or non-sink-like anchors can distort future-fate estimates. Robust conclusions should remain qualitatively stable across plausible horizon and anchor settings, while low selected-path coverage should limit confidence in projected vectors and Signed Ordering Flux. For condition comparisons, the biological replicate but not the individual cell, must remain the unit of inference. Large cell numbers cannot compensate for too few independent donors, animals, organoids or cultures, and rebuilding separate graphs for each condition may introduce structural differences that are unrelated to the biological effect of interest. scCS therefore uses a pooled scientific model by default and treats condition- or replicate-blocked graphs as sensitivity analyses. Likewise, downstream gene associations and enrichment results are intended to nominate candidate mechanisms, not to establish causality.

Within these boundaries, scCS provides a transparent way to quantify a clearly stated fate hypothesis without replacing or redefining the source dynamics.

## Methods

### Biological and mathematical foundations of scCS

Single-cell transcriptomic measurements provide molecular snapshots of individual cells rather than direct observations of their complete developmental histories or eventual outcomes. Trajectory-inference methods address this limitation by organizing cells along continuous or branching progression coordinates, including pseudotime and diffusion-based orderings[20,21]. RNA velocity adds directional information by estimating the local derivative of transcriptional state from spliced and unspliced RNA abundances[3,4]. These quantities are informative about progression and short-term state change, but they do not directly observe the terminal fate ultimately reached by an individual cell.

Cell-fate determination is not necessarily represented by a single deterministic path through transcriptional space. Differentiating cells can occupy continuous intermediate states, retain support for multiple possible outcomes and exhibit fate biases before complete molecular separation of terminal populations. Experimental lineage tracing has shown that transcriptional state can contain predictive information about future fate while still failing to determine fate uniquely, including cases in which fate-relevant differences are not fully visible in the measured transcriptome[14]. Probabilistic fate-mapping approaches consequently represent differentiation as a stochastic process and assign distributions over possible terminal outcomes rather than a single mandatory trajectory[5,6].

This biological distinction motivates the separation of local direction, future outcome support and fate resolution in scCS. RNA velocity or another directed transition model describes the distribution of plausible immediate movements from each measured state. Future-fate probabilities instead summarize the cumulative consequences of repeatedly following those transitions. A cell can therefore have a local vector directed away from a fate while retaining future graph paths that later reach that fate, or it can point transiently toward a fate while having little total probability of reaching the corresponding terminal population. Instantaneous direction and finite-horizon future identity are thus related but non-identical estimands.

scCS represents the source dynamics as a row-stochastic transition matrix P, where Pij is the probability assigned to a one-step transition from cell i to cell j. This Markov representation assumes that the next-state distribution is conditioned on the currently represented cell state and transition model. Similar directed Markov formulations have been used to calculate fate probabilities and terminal-state absorption in single-cell analysis[6,12]. scCS differs in that the root population and candidate outcomes are supplied as a supervised biological hypothesis rather than necessarily discovered from the transition matrix. The graph provides dynamical evidence against which that hypothesis is evaluated.

The scCS future-fate formulation combines absorbing outcome states with discounted propagation. Cells selected as endpoint anchors represent the supplied outcomes and are made absorbing. From any non-anchor cell, the process follows the transition graph while it continues, but at every step it can stop without reaching an anchor. This geometric stopping mechanism introduces an explicit finite expected graph depth and retains probability that remains unresolved. The absorbing-state component is related to established Markov-chain fate mapping[6], whereas the combination of user-supplied anchors, finite-horizon discounting and an explicit unresolved outcome defines the DFFP formulation introduced in scCS.

The resulting outcome probabilities are not treated as a single commitment score. scCS separates the total probability reaching any selected fate from the conditional distribution among those fates. Entropy of the conditional distribution measures how specifically the resolved future favors one candidate outcome, whereas unresolved probability records the future mass not assigned to any modeled anchor at the declared horizon. Signed Ordering Flux is computed independently from expected one-step change along a supplied progression coordinate. This decomposition distinguishes whether the modeled fates are reachable, which fate is favored, how decisive that preference is and whether the measured local dynamics are moving forward or backward along the biological ordering.

### Software design and data model

scCS is implemented in Python and operates on AnnData objects. The core dependencies are NumPy, SciPy, pandas, matplotlib, AnnData and Scanpy[22]; scVelo[4] is an optional dependency for RNA-velocity estimation and graph extraction, statsmodels is optional for gene-association and mixed-model procedures, and gseapy is optional for remote enrichment. The principal public interfaces are SingleScorer, PairScorer and MultiScorer. Lower-level functions expose transition canonicalization, ordering scaling, supervised embedding, DFFP solution, instantaneous projection, population summaries, inference and downstream analyses.

The software accepts a precomputed transition matrix rather than requiring a single velocity estimator. A typical workflow obtains neighbors, moments, velocity and a velocity graph with scVelo, but any non-negative directed matrix with rows interpretable as transition probabilities can be supplied after canonicalization. scCS stores fitted metrics and provenance in AnnData, can export tables and metadata as CSV or JSON, and can serialize fitted scorer objects for reproducible reuse.

### Scientific inputs and furcation specifications

A supervised furcation consists of (i) one root label; (ii) K≥2 candidate fate labels; (iii) a categorical annotation field containing those labels; (iv) a continuous ordering value for each cell; and (v) a directed transition matrix over the full set of cells. Optional inputs include explicit competing outcomes, condition labels, biological-replicate identifiers and covariates for downstream association analysis. The selected set S contains cells in the supplied root and selected fates. This set defines scoring summaries and display positions, not the state space of DFFP. Cells outside S remain in the transition matrix and can mediate future paths. Explicit competing outcomes are biologically named absorbing outcomes that are included in the solve but not included in selected-fate CFA, DFR, FFS or RC. Probability not assigned to either selected or explicit competing outcomes before stopping is UFP.

### Preflight validation

Preflight checks are performed before fitting. The checks verify that annotation, ordering, condition and replicate columns exist; root and fate labels are present with sufficient cells; ordering values are finite; the transition matrix has the expected square shape; and required velocity metadata are available when package-managed velocity estimation is requested. Condition analyses additionally verify that replicate identifiers are not confounded with missing condition labels and that the requested groups contain biological replicates. Diagnostics are returned as structured error, warning and information records and can be converted to a table or raised as exceptions.

### Transition-matrix canonicalization

Let P∈R^{N×N} denote the supplied transition matrix. Dense or sparse inputs are converted to compressed sparse row form. Small negative numerical values are clipped to zero; materially negative entries are rejected. Each nonzero row is normalized to sum to one. A zero row receives a self-loop so that every row is stochastic. The canonical matrix therefore satisfies Pij≥0 and ΣjPij=1. Canonicalization metadata record the original format, row-sum behavior and any corrections.

### Ordering extraction, orientation and scaling

The continuous ordering may be latent time, velocity pseudotime, diffusion pseudotime, developmental potential transformed to the requested direction, or experimentally measured time. The user declares whether higher values represent later states. scCS supports rank scaling, min–max scaling or no scaling. Rank scaling is robust to nonlinear spacing and maps valid values to a common interval; min–max scaling preserves relative distances but is more sensitive to outliers. Root and fate distributions are summarized to diagnose reversed or weakly separated orderings.

The ordering has three roles that remain explicit: it positions cells along display rays, identifies late cells within each annotated fate for endpoint anchors, and defines the sign of local progression. It does not determine graph transitions and does not by itself generate fate probabilities.

### Standardized supervised geometry

For K fates, scCS constructs K unit fate directions as vertices of a regular simplex. The directions satisfy equal pairwise angles:

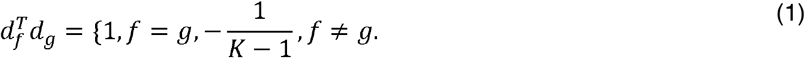

An incoming root axis is placed orthogonal to the fate simplex. Root cells are positioned along this incoming axis and fate cells along their corresponding outgoing direction according to the scaled ordering. Deterministic jitter can be added for display. This embedding is a scientific comparison coordinate system and a visualization; DFFP probabilities are never computed by rebuilding neighbors in this space.

### Endpoint-anchor construction

Within each fate annotation, endpoint anchors are selected from cells at or above a specified ordering quantile. A minimum anchor count prevents extremely small absorbing sets; when necessary, the latest cells are added deterministically to reach the minimum. Optional competing outcomes are anchored by the same logic or supplied explicit masks. A cell belongs to at most one outcome anchor set.

Anchor diagnostics quantify, for each anchor set, outgoing mass to the same fate, the root, other selected fates and cells outside S, together with mean ordering and the fraction of transitions that move forward. Anchors are not required to be perfect sinks. Non-sink-like anchors are reported because they indicate that the biological annotation and the graph dynamics are not fully aligned.

### Discounted Future-Fate Propagation

DFFP implements the finite-horizon absorbing Markov formulation introduced above. Anchor rows are made absorbing to form P*. Let *A*∈*{0,1}*^*N×O*^ (where *H* ⋅*=A, i* ∈ *A*) identify O selected and optional competing outcomes. Before each graph transition, a process continues with probability γ and stops unresolved with probability 1−γ. The effective horizon h is defined by

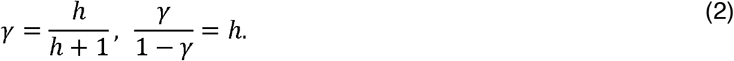

Thus, h is the expected number of continued transitions under geometric stopping. It specifies graph depth and is not interpreted as physical time. For an anchor cell, the outcome vector equals its anchor indicator. For a transient cell i, the discounted outcome vector Hi satisfies

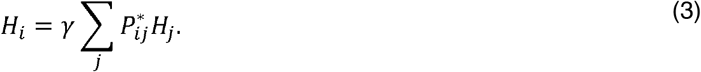

After ordering transient cells before anchor cells, write Q for the transient-to-transient block and R for the transient-to-anchor block. The transient outcome matrix HT is the solution of

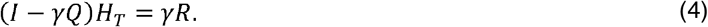

For graphs up to the configured direct-solver threshold, scCS factorizes the sparse matrix and solves all outcome right-hand sides. For larger graphs, it uses the sparse fixed-point update

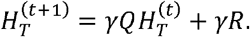

Iteration continues until the maximum update and linear-system residual satisfy tolerance or the maximum iteration count is reached. Convergence status, iteration count and residual are retained. The unresolved probability is

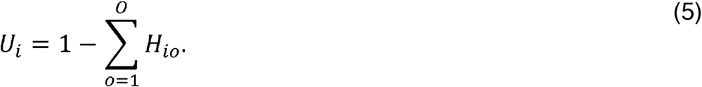

Numerical values are clipped only within a small tolerance and probability-conservation checks are reported. Anchor cells have U=0 for their assigned outcome. Because the solve uses the full P*, paths can traverse cells outside the selected furcation and later return. This differs from subsetting and renormalizing the transition matrix before the solve.

Unresolved Future Probability (UFP), defined in equation (5), is the probability not assigned to any selected or explicit competing anchor before geometric stopping. The absolute contribution of selected fate f is pif = Ri qif, which is identical to the DFFP outcome probability for that fate. Thus, DFR reports how much of the future reaches the selected outcomes, CFA reports how that reached mass is distributed, FFS reports how decisive that distribution is, RC combines reach and specificity, and UFP reports what remains unresolved. High CFA with low DFR should not be interpreted as strong commitment, and hard dominant-fate or status labels are secondary summaries of the continuous quantities.

### Selected-fate metrics

Let pif denote the DFFP probability for selected fate f∈{1,…,K}. Discounted Fate Reach is the total probability assigned to selected fates:

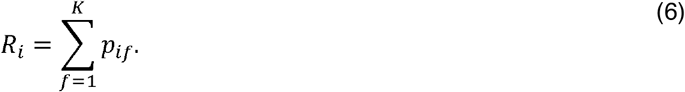

When Ri is at least the configured minimum reach, Conditional Fate Affinity is

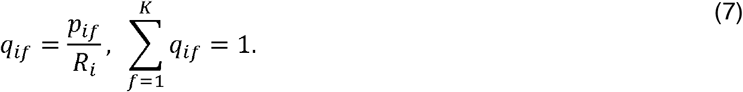

For cells below the minimum reach, scCS uses a neutral uniform vector for numerical completeness and marks the cell as low reach; such CFA values should not be interpreted as resolved identity. Normalized conditional entropy Ei and Future-Fate Specificity Si are

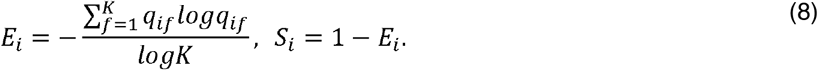

Terms with *q*_*if*_ = 0 are evaluated using the convention 0 log0 = 0.

Resolved Commitment combines selected-fate reach with conditional specificity:

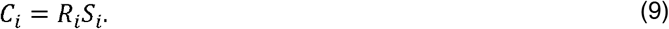

The absolute contribution of fate f to the selected future is pif=Riqif. These components are retained so that a composite RC cannot obscure whether a low value arises from poor reach or ambiguity. Dominant fate, angular summaries, entropy and status categories are secondary outputs derived from the continuous quantities.

### Signed Ordering Flux and selected-path coverage

For cell i, one-step transition mass retained within the selected set S is

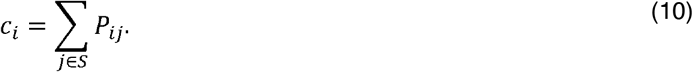

When ci>0, transitions within S are conditioned as

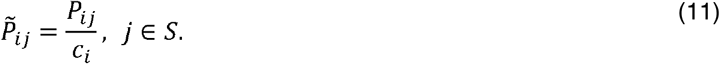

Signed Ordering Flux is the expected one-step change in the supplied ordering:

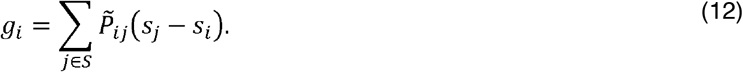

Positive values indicate forward local movement, negative values indicate retrograde movement and values near zero can arise from stationary, balanced, turning or loop-like transitions. The support-weighted flux cig_i is also returned. Coverage ci is always reported because a signed flux estimated from little retained transition mass describes only a restricted part of the source transition distribution.

### Transition-mediated projection into the supervised geometry

scCS does not directly rotate or warp velocity vectors from an unsupervised embedding into the supervised star. Instead, the directed cell-cell transition operator provides a common representation of the dynamics. The same transition probabilities that describe movement between cells in the source manifold are applied to coordinates assigned in the supervised geometry, thereby transferring dynamical relationships rather than the Euclidean orientation of arrows in a particular embedding.

For a selected furcation containing the cell set S, let Pij denote the source transition weight from cell i to cell j. The retained outgoing mass, total outgoing mass and selected-path coverage are

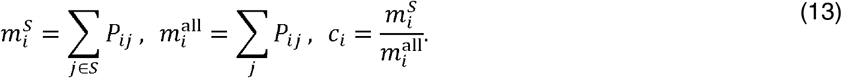

Transitions to cells outside S are not assigned artificial star coordinates. Their probability remains external transition mass and is retained as a diagnostic. When the retained mass is positive, the within-furcation weights are conditionally normalized as

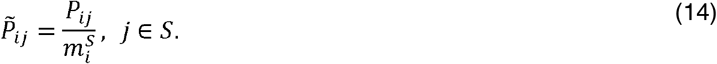

For standardized scientific coordinates yi, the transition-induced vector is the transition-weighted expected displacement from the source cell to its possible destinations:

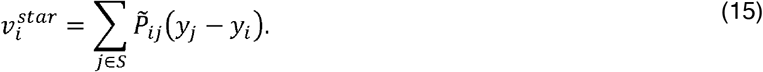

Equivalently, with coordinate matrix Y and row-normalized retained transition matrix P-tilde, the projected field is

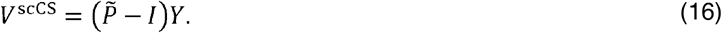

This operation is a graph-based analogue of a vector-field pushforward, but it requires neither a differentiable mapping nor an inverse transformation between embeddings. Crucially, scCS does not rebuild neighbors in the supervised geometry. Cells with no retained outgoing mass, or coverage below a declared threshold, receive an undefined or suppressed projected vector rather than a potentially misleading direction.

The projected vector is separated into motion along the root-progression axis r and motion within the terminal-fate subspace:

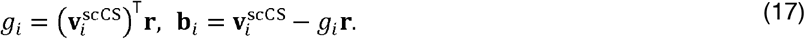

Here gi is the signed local progression component and bi is the fate-directed component. Positive gi indicates movement in the forward orientation of the supplied ordering, negative values indicate retrograde motion and values near zero can arise from stationary, balanced, turning or loop-like transitions.

The fate-directed component is compared with the regular-simplex directions. If theta_if is the angle between bi and fate direction df, cosine-softmax affinity is

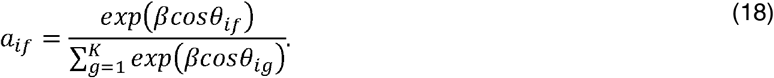

The inverse-temperature parameter beta is calibrated from a requested probability for a perfectly aligned direction under the simplex geometry. Direction magnitude can be transformed with a robust Hill function so that a small number of large vectors do not dominate visualization. Instantaneous affinity is a local directional quantity and is not interpreted as DFFP, a hitting probability or a long-term fate probability.

For the two-dimensional display star, transition-induced vectors are recomputed from the same normalized transition weights and the displayed coordinates xi:

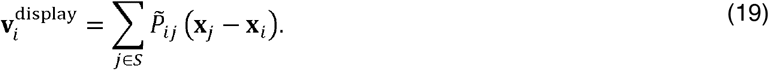

The display vector is therefore not obtained by projecting the K-dimensional scientific vector onto two arbitrary axes. Recalculation from the transition operator ensures that an arrow represents expected movement between the cell locations shown in the display. Two-dimensional velocity grids, rose plots and branch-relative profiles are visualization and quality-control products only; they do not modify the scientific scores, and DFFP remains solved on the original full transition graph.

### Estimand-based method selection and sensitivity analysis

The scoring mode is selected before inspecting the preferred visual result. Future-fate mode is used when the target is discounted future-fate support at the declared effective horizon for supplied outcomes, including unresolved mass and paths outside S. Instantaneous mode is used when the target is the direction of immediate transition-induced motion in the supervised geometry. First-exit or long-time absorption calculations can be used as explicit sensitivity analyses when those alternative estimands are scientifically relevant, but they are not substituted for DFFP without changing the interpretation.

DFFP sensitivity analysis varies effective horizon, anchor quantile, minimum anchor count and, when justified, explicit competing outcomes. Results should report whether root and population CFA, DFR, FFS, RC and SOF are qualitatively stable; whether solver residuals remain acceptable; and whether anchor diagnostics change. Sensitivity is evaluated on aligned cell identifiers when comparing alternative velocity models or graph constructions.

### Population summaries and status composition

Population summaries calculate means, medians, dispersion and distributions for root and fate annotations. Root future composition aggregates selected-fate contributions and normalizes them to the total selected-fate mass. Pairwise population contrasts and subset summaries are available. Status-composition plots report the proportion of cells classified as unresolved, ambiguous, low reach or fate committed according to declared thresholds. Population balance entropy is calculated from aggregated composition and is not equated with mean cell-level FFS.

### Visualization and data products

SingleScorer provides star maps, projected velocity displays, direction-strength maps, rose plots, population commitment plots, branch velocity profiles, pairwise summaries, bar plots, heatmaps and multi-panel summaries. The native manifold remains the primary location for assessing the velocity field. The standardized star is used to compare supplied branches and fitted quantities on a common geometry. Plotting can subsample cells for rendering, but scoring and inference operate on the full selected data.

Fitted quantities can be written to AnnData observation columns and multidimensional arrays, exported as tables, or retained in result dataclasses. Provenance includes scorer mode, fate names, ordering configuration, transition scope, DFFP parameters, anchor metadata and solver diagnostics. Scorer save/load methods preserve the fitted analysis for later visualization and downstream analysis.

### Condition-level model construction

ConditionScorer provides shared infrastructure for PairScorer and MultiScorer. The default analysis constructs one pooled supervised embedding and one pooled transition-based scientific model across all conditions, then computes cell-level metrics and aggregates them within biological replicates. This makes replicate effects comparable under the same coordinate system, anchors and graph-based estimand.

The default transition scope is pooled. Condition-blocked and replicate-blocked graphs are available as sensitivity analyses because blocking prevents transitions across groups and may remove biologically valid manifold neighbors. The chosen scope and its graph-coverage consequences are included in the output. The package does not treat cell labels as exchangeable experimental units.

### Construction of pseudo-conditions for MultiScorer validation

The control, low and high groups in the Schwann-cell analysis were constructed solely to validate the MultiScorer workflow and do not represent biological treatment conditions. To obtain approximately balanced pseudo-conditions and pseudo-replicates, cells were stratified jointly by their annotated population and by up to five quantile bins of inverse CytoTRACE2 score. Cell identifiers were shuffled independently within each stratum using random seed 20260714 and assigned cyclically among 12 condition–replicate combinations comprising control, low and high conditions with four pseudo-replicates per condition. This procedure preserved the distribution of annotated states and progression values across the constructed groups. The resulting conditions contained 2,965, 2,940 and 2,916 cells, respectively. Within the modeled root population, control contained 235 cells and low and high each contained 220 cells, corresponding to 55–60 root cells per pseudo-replicate.

A controlled condition effect was introduced by reweighting outgoing transitions from Common Progenitor cells toward destinations with greater baseline support for the ChC fate. A baseline SingleScorer model was first fitted to the original dynamical RNA-velocity transition matrix using an effective horizon of 64, an anchor quantile of 0.90, a minimum of 10 anchor cells and rank-scaled progression. Let s_j ∈ [0,1] denote the baseline Conditional Fate Affinity toward ChC for destination cell j, with annotated ChC cells assigned s_j = 1. For a root cell i belonging to condition c(i) and pseudo-replicate r(i), each existing outgoing transition was reweighted as follows:

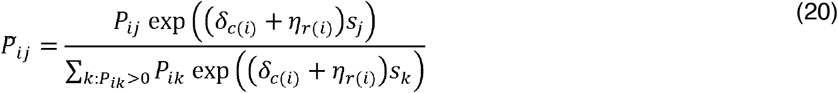

where the condition-level log shifts were

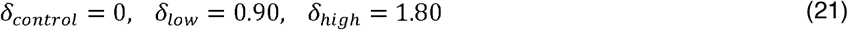

and pseudo-replicate effects were sampled independently as

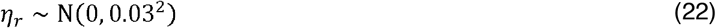

using random seed 20260715. Only transition weights from root cells were modified; cell annotations, expression values, progression coordinates, non-root transition rows and the set of nonzero graph edges were unchanged. Each modified row was renormalized to sum to one, preserving a row-stochastic transition matrix and the original graph support.

MultiScorer was then fitted once in future-fate mode using a pooled transition graph, shared endpoint anchors and a common supervised geometry across the three pseudo-conditions. Condition summaries were calculated for the root population, requiring at least 20 cells and all four pseudo-replicates per condition. Omnibus comparisons used one-way replicate-label permutation tests with 9,999 Monte Carlo permutations. Adjusted omnibus P values were calculated using the Holm procedure across the tested fates for each metric. Pairwise ChC comparisons used exact replicate-label permutation tests; with four pseudo-replicates in each group, all 70 possible assignments were enumerated for each two-condition comparison, followed by Holm adjustment across the three pairwise tests. The prespecified ordered contrast used weights −1, 0 and 1 for control, low and high, respectively, and was evaluated using 9,999 permutations. Because the conditions and replicate labels were constructed for software validation, the resulting effect estimates and P values demonstrate recovery and reporting behavior rather than biological evidence.

### Pairwise replicate-level inference

For two independent conditions A and B, let zr denote a replicate-level summary of the requested metric and let G(r) denote its condition. The estimated effect is

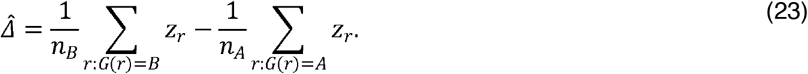

The metric can be total DFR, FFS, RC, SOF or fate-specific CFA or contribution. When the number of possible replicate-label assignments is below a configured threshold, all assignments with the observed group sizes are enumerated. Otherwise, Monte Carlo permutations are sampled. For B random permutations and b statistics at least as extreme as the observed statistic, the two-sided finite-sample P value uses (b+1)/(B+1). Multiple outcomes can be adjusted with Benjamini–Hochberg or another declared family-wise procedure.

Hierarchical bootstrap confidence intervals first sample biological replicates with replacement within condition and then sample cells with replacement within each selected replicate[23]. This retains both between-replicate and within-replicate variability. Delta-CS and trajectory-shift summaries decompose changes across fates and progression metrics, and plotting functions expose replicate values rather than only condition means.

### Multi-condition inference, post-hoc tests and contrasts

For three or more independent conditions, MultiScorer calculates a one-way F statistic from replicate summaries and permutes condition labels while preserving group sizes. A significant omnibus result can be followed by pairwise replicate-label permutation tests with Holm adjustment. Planned contrasts are specified by weights that sum to zero and are tested against the corresponding replicate-label permutation distribution. Omnibus, post-hoc and contrast analyses retain the metric, fate, number of replicates, permutation mode and adjustment metadata.

### Mixed-model sensitivity analysis

An optional random-intercept linear mixed model can be fitted to cell-level or summarized outcomes with biological replicate as the grouping factor. The implementation audits optimizer convergence, random-effect variance, fixed-effect covariance and singularity. Invalid or degenerate fits return an explicit failure audit rather than a coefficient that appears inferentially valid. Mixed models are supplementary to the replicate-permutation analysis and do not repair a design with too few independent replicates.

### Expression visualization and trends

Measured expression can be displayed on the standardized star without changing cell positions or fitted scores. Expression trends are calculated by binning cells along the supplied ordering or a selected fitted quantity such as CFA, fate contribution or RC and plotting bin summaries. These curves are descriptive because neighboring cells and bins are not independent experimental units. Shared color scales are supported for direct condition comparison; gene-specific scales can be used when dynamic ranges differ.

### Commitment-associated genes

get_commitment_associated_genes tests partial rank association between expression and a selected fitted outcome. Expression and outcome variables are converted to ranks. Covariates, including the supplied ordering, are residualized from both ranked variables by linear projection, and the association is the correlation of residuals. Sparse matrices are processed in gene chunks. P values are adjusted with the Benjamini–Hochberg procedure.

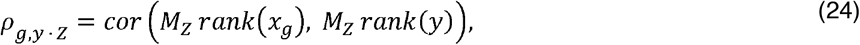

where MZ is the residual-maker for the covariate matrix Z. Cell-level mode is labeled exploratory. When genuine replicate metadata are available, expression and outcomes can be aggregated within replicates before formal association. The method ranks candidate genes associated with the fitted quantity; it does not establish causal lineage regulation.

### Fate markers and enrichment

get_fate_markers compares each annotated fate population with the root to identify identity markers. This analysis is distinct from association with variation in commitment among root cells. Enrichment analysis accepts a local gene-set mapping or GMT file and performs over-representation analysis with Fisher’s exact test followed by multiple-testing correction. A user-specified assay-relevant background is preferred. Remote Enrichr access is optional[24] and should be accompanied by library name and access date because remote libraries can change.

### Example datasets

The pancreas example used a processed mouse pancreatic endocrinogenesis dataset[13] distributed with scVelo[4]. Neighbors, moments, velocity, a velocity graph and a continuous ordering were calculated before scCS. Ngn3-high endocrine progenitor and pre-endocrine cells were treated as the root population, and Alpha, Beta, Delta and Epsilon populations as candidate fates. The example demonstrates a four-fate furcation and branches with forward, turning and retrograde local dynamics.

The neural crest–Schwann-cell example used the Schwann dataset distributed with RegVelo[11]. Common Progenitor was supplied as root and Gut, Gut neuron and chromaffin cells as candidate fates. Dynamical and deterministic velocity models can be compared while preserving the same cells and graph context. The example demonstrates a three-fate problem with lower root reach and a strongly retrograde annotated branch.

Control, low and high pseudo-conditions were generated as controlled software-validation labels using the procedure described under “Construction of pseudo-conditions for MultiScorer validation.” They are not biological treatment groups, and their effect estimates and *P*values are not interpreted as biological discoveries.

### Scalability and numerical benchmarking

Two workloads are benchmarked separately. After outcome probabilities have been solved, calculation of CFA, DFR, FFS, RC and UFP is cellwise with memory proportional to N×K. The full DFFP solve stores the sparse transition matrix and one or more N×O state-by-outcome arrays, with memory scaling approximately as the number of retained graph edges plus N×O. Direct and iterative solvers have different computational regimes.

No-chunk benchmarks allocate each complete problem in memory. A target is labeled measured only if the full graph and solver state were allocated and the solve completed under the stated tolerance. Targets that exceed the machine’s safety fraction are labeled skipped rather than replaced by streaming. Analytic memory estimates and runtime extrapolations are reported separately from measured runs. Neighbor construction, kinetic fitting and velocity-graph estimation are upstream procedures and are excluded from scCS-only throughput claims.

### Software testing and reproducibility

Unit tests cover transition canonicalization, ordering, furcation geometry, instantaneous affinity, DFFP probability conservation and solver behavior, population summaries, condition inference, mixed-model audits, downstream gene and enrichment functions, scorer pipelines and AnnData integration. The supplied source snapshot includes twelve tutorial notebooks spanning single-dataset, pairwise, multi-condition, method-selection, complex-branch, downstream and scalability workflows. The archived analysis records the package release, Python and dependency versions, random seeds, graph construction, ordering field, root and fate annotations, condition and replicate columns, DFFP parameters, solver tolerance and sensitivity settings.

## Data availability

The example datasets are available through the upstream scVelo and RegVelo software resources and their source publications[11,13]. No new primary biological data were generated for this software study.

## Code availability

scCS source code is available at https://github.com/mcrewcow/scCS and documentation at https://sccs-py.readthedocs.io/. scCS is available on PyPi at https://pypi.org/project/scCS-py/. The manuscript was prepared from the 0.8.0 source snapshot.

## Analysis reproducibility

The results presented in this manuscript were prepared using the following setup: 2 x Intel Xeon w9 (2 x 56 cores)

1.1 TB RAM

Ubuntu 24.04 LTS

Python 3.12.13

Scalability benchmarks were run using 1 core.

## Acknowledgements

This work was supported by the grants from the Gilbert Family Foundation (PB) and Department of Defense VRP FTTSA VR220053 (PB). The authors express gratitude to the research community for the publicly available data, used in this study.

## Author contributions

E.K. conceived and developed scCS, designed and implemented the methodology and software, performed the analyses, prepared the figures and wrote the original manuscript. E.K. and E.I. tested the software. E.K. validated the software. E.K, P.B. wrote the manuscript. E.K., E.I., P.B. edited the manuscript. P.B. supervised the research. All authors reviewed and approved the final manuscript.

## References

[1] Street K, Risso D, Fletcher RB, et al. Slingshot: cell lineage and pseudotime inference for single-cell transcriptomics. BMC Genom 2018;19:477. 10.1186/s12864-018-4772-0.

[2] Cao J, Spielmann M, Qiu X, et al. The single-cell transcriptional landscape of mammalian organogenesis. Nature 2019;566:496–502. 10.1038/s41586-019-0969-x.

[3] Manno GL, Soldatov R, Zeisel A, et al. RNA velocity of single cells. Nature 2018;560:494–8. 10.1038/s41586-018-0414-6.

[4] Bergen V, Lange M, Peidli S, et al. Generalizing RNA velocity to transient cell states through dynamical modeling. Nat Biotechnol 2020;38:1408–14. 10.1038/s41587-020-0591-3.

[5] Setty M, Kiseliovas V, Levine J, et al. Characterization of cell fate probabilities in single-cell data with Palantir. Nat Biotechnol 2019;37:451–60. 10.1038/s41587-019-0068-4.

[6] Lange M, Bergen V, Klein M, et al. CellRank for directed single-cell fate mapping. Nat Methods 2022;19:159–70. 10.1038/s41592-021-01346-6.

[7] Steinschaden T, Faure L, Soldatov R, et al. A competition model of multilineage priming and cell-fate decisions. Cell Rep 2026;45:116690. 10.1016/j.celrep.2025.116690.

[8] Squair JW, Gautier M, Kathe C, et al. Confronting false discoveries in single-cell differential expression. Nat Commun 2021;12:5692. 10.1038/s41467-021-25960-2.

[9] Kang M, Gulati GS, Brown EL, et al. Improved reconstruction of single-cell developmental potential with CytoTRACE 2. Nat Methods 2025;22:2258–63. 10.1038/s41592-025-02857-2.

[10] Li C, Virgilio MC, Collins KL, et al. Multi-omic single-cell velocity models epigenome– transcriptome interactions and improves cell fate prediction. Nat Biotechnol 2022:1–12. 10.1038/s41587-022-01476-y.

[11] Wang W, Hu Z, Weiler P, et al. RegVelo: Gene-regulatory-informed dynamics of single cells. Cell 2026. 10.1016/j.cell.2026.04.022.

[12] Weiler P, Lange M, Klein M, et al. CellRank 2: unified fate mapping in multiview single-cell data. Nat Methods 2024;21:1196–205. 10.1038/s41592-024-02303-9.

[13] Bastidas-Ponce A, Tritschler S, Dony L, et al. Comprehensive single cell mRNA profiling reveals a detailed roadmap for pancreatic endocrinogenesis. Development 2019;146:dev173849. 10.1242/dev.173849.

[14] Weinreb C, Rodriguez-Fraticelli A, Camargo FD, et al. Lineage tracing on transcriptional landscapes links state to fate during differentiation. Science 2020;367:eaaw3381. 10.1126/science.aaw3381.

[15] Keren-Shaul H, Spinrad A, Weiner A, et al. A Unique Microglia Type Associated with Restricting Development of Alzheimer’s Disease. Cell 2017;169:1276–1290.e17. 10.1016/j.cell.2017.05.018.

[16] Krasemann S, Madore C, Cialic R, et al. The TREM2-APOE Pathway Drives the Transcriptional Phenotype of Dysfunctional Microglia in Neurodegenerative Diseases. Immunity 2017;47:566–581.e9. 10.1016/j.immuni.2017.08.008.

[17] Kriukov E, Mukwaya A, Cullen PF, et al. Single-cell transcriptome of retinal myeloid cells in response to transplantation of human neurons reveals reversibility of microglial activation. bioRxiv 2025:2025.05.16.654622. 10.1101/2025.05.16.654622.

[18] Srivatsan SR, McFaline-Figueroa JL, Ramani V, et al. Massively multiplex chemical transcriptomics at single-cell resolution. Science 2020;367:45–51. 10.1126/science.aax6234.

[19] Sande BV de, Lee JS, Mutasa-Gottgens E, et al. Applications of single-cell RNA sequencing in drug discovery and development. Nat Rev Drug Discov 2023;22:496–520. 10.1038/s41573-023-00688-4.

[20] Trapnell C, Cacchiarelli D, Grimsby J, et al. The dynamics and regulators of cell fate decisions are revealed by pseudotemporal ordering of single cells. Nat Biotechnol 2014;32:381–6. 10.1038/nbt.2859.

[21] Haghverdi L, Büttner M, Wolf FA, et al. Diffusion pseudotime robustly reconstructs lineage branching. Nat Methods 2016;13:845–8. 10.1038/nmeth.3971.

[22] Wolf FA, Angerer P, Theis FJ. SCANPY: large-scale single-cell gene expression data analysis. Genome Biol 2018;19:15. 10.1186/s13059-017-1382-0.

[23] Saravanan V, Berman GJ, Sober SJ. Application of the hierarchical bootstrap to multi-level data in neuroscience. Neurons, Behav, Data Anal Theory 2020;

[24] Kuleshov MV, Jones MR, Rouillard AD, et al. Enrichr: a comprehensive gene set enrichment analysis web server 2016 update. Nucleic Acids Res 2016;44:W90–7. 10.1093/nar/gkw377.

