## Supplementary figures and images for "Quantitative assessment of cell fate commitment in single-cell transcriptomics using scCS"

### Figure 1

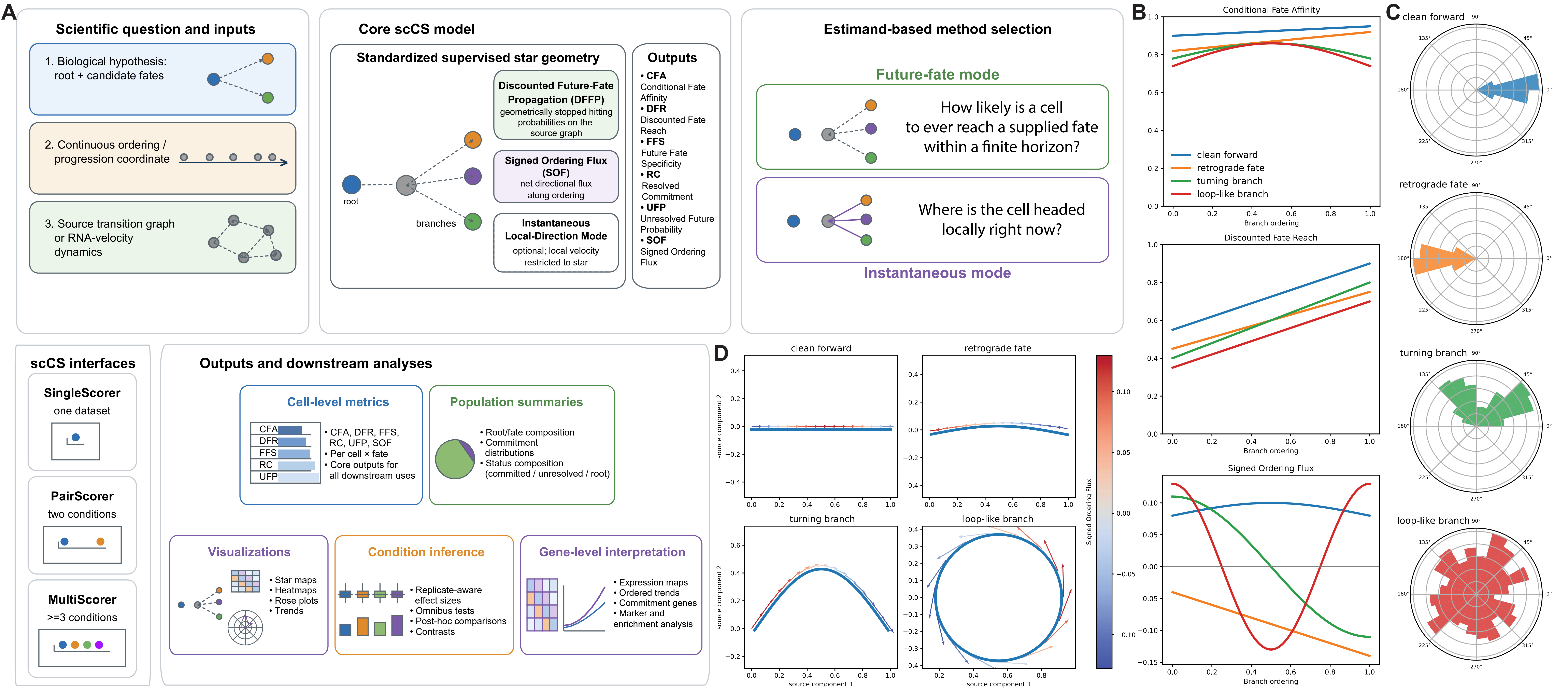

### Figure 2

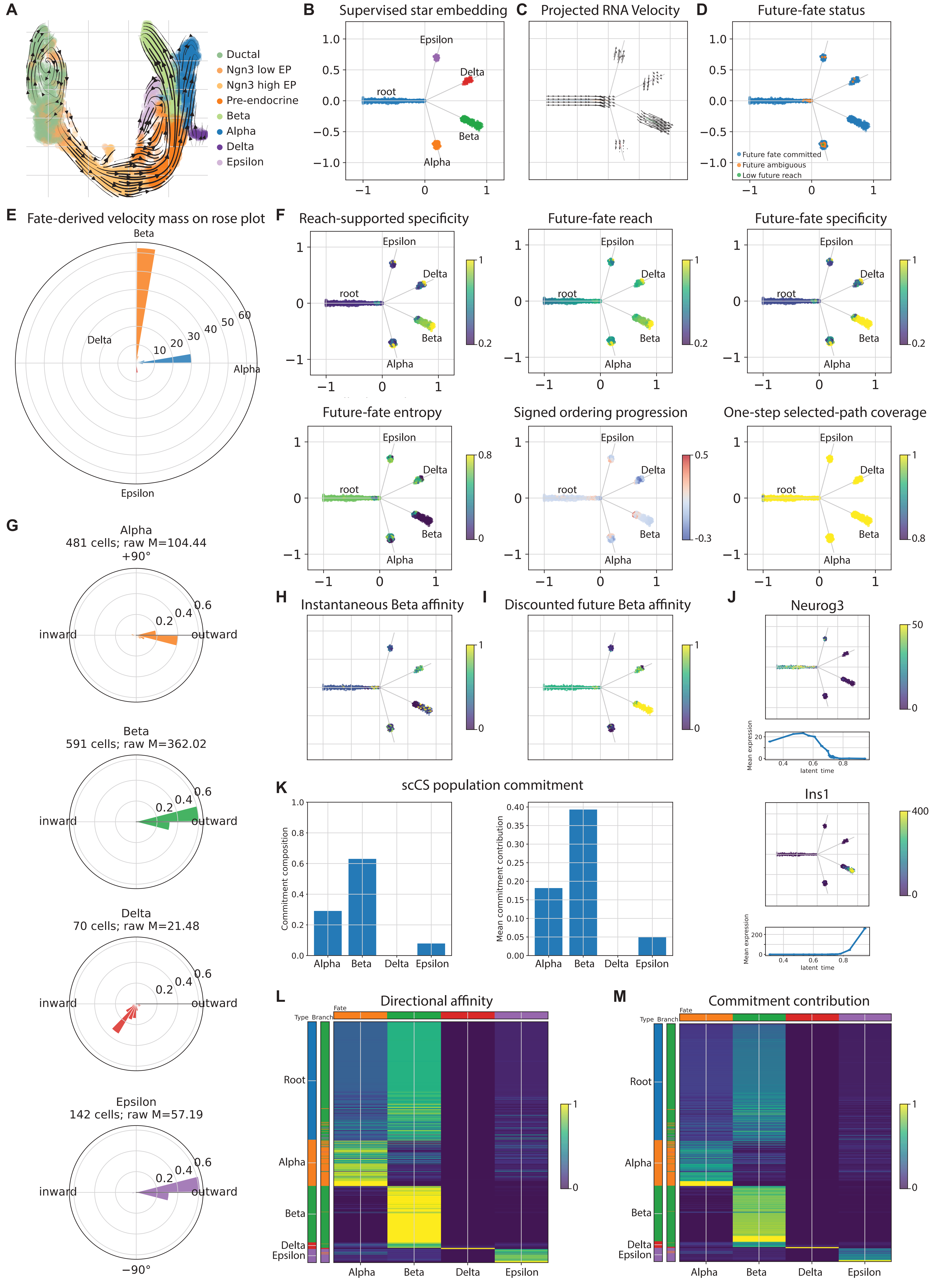

### Figure 3

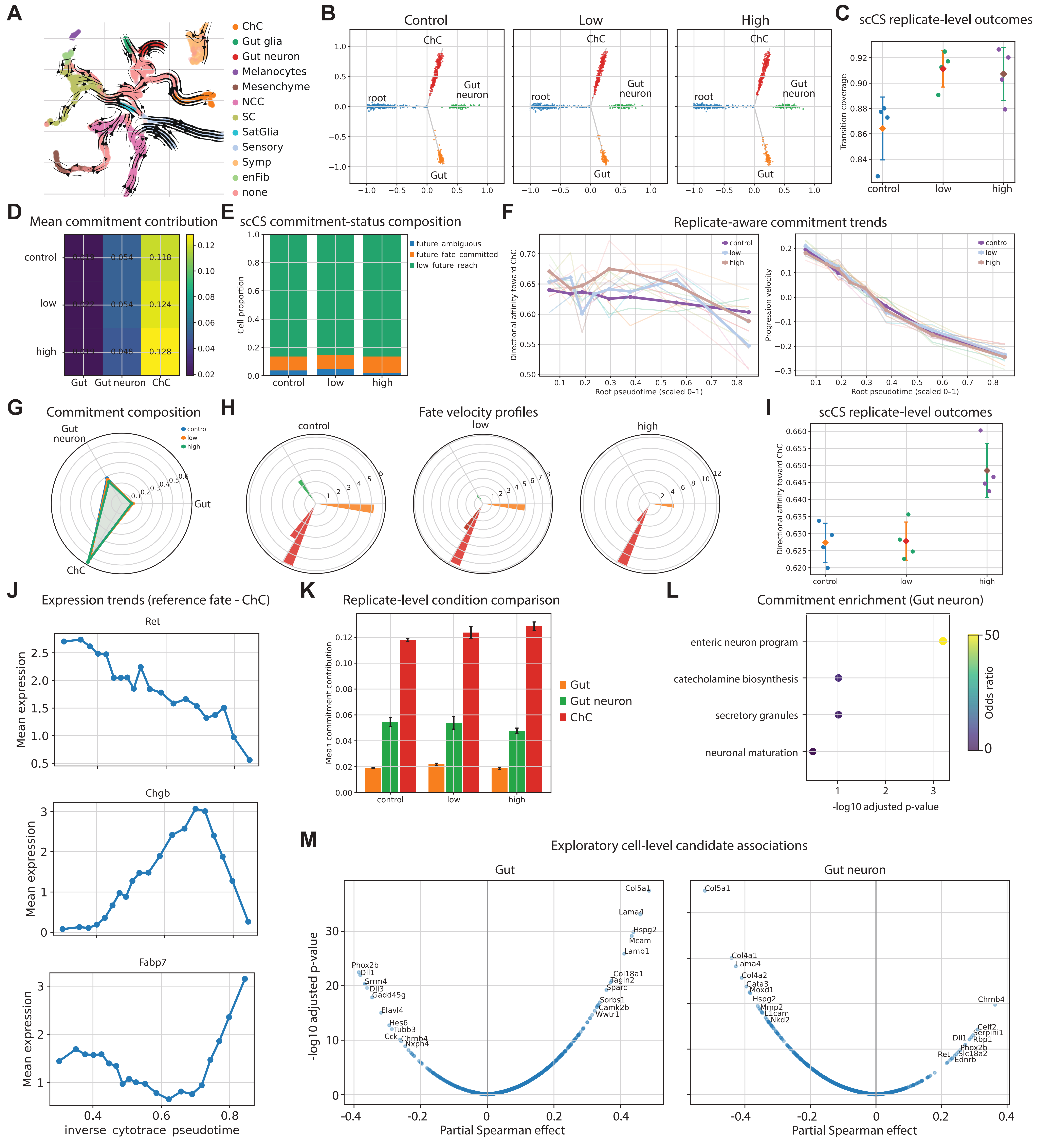

### Figure 4

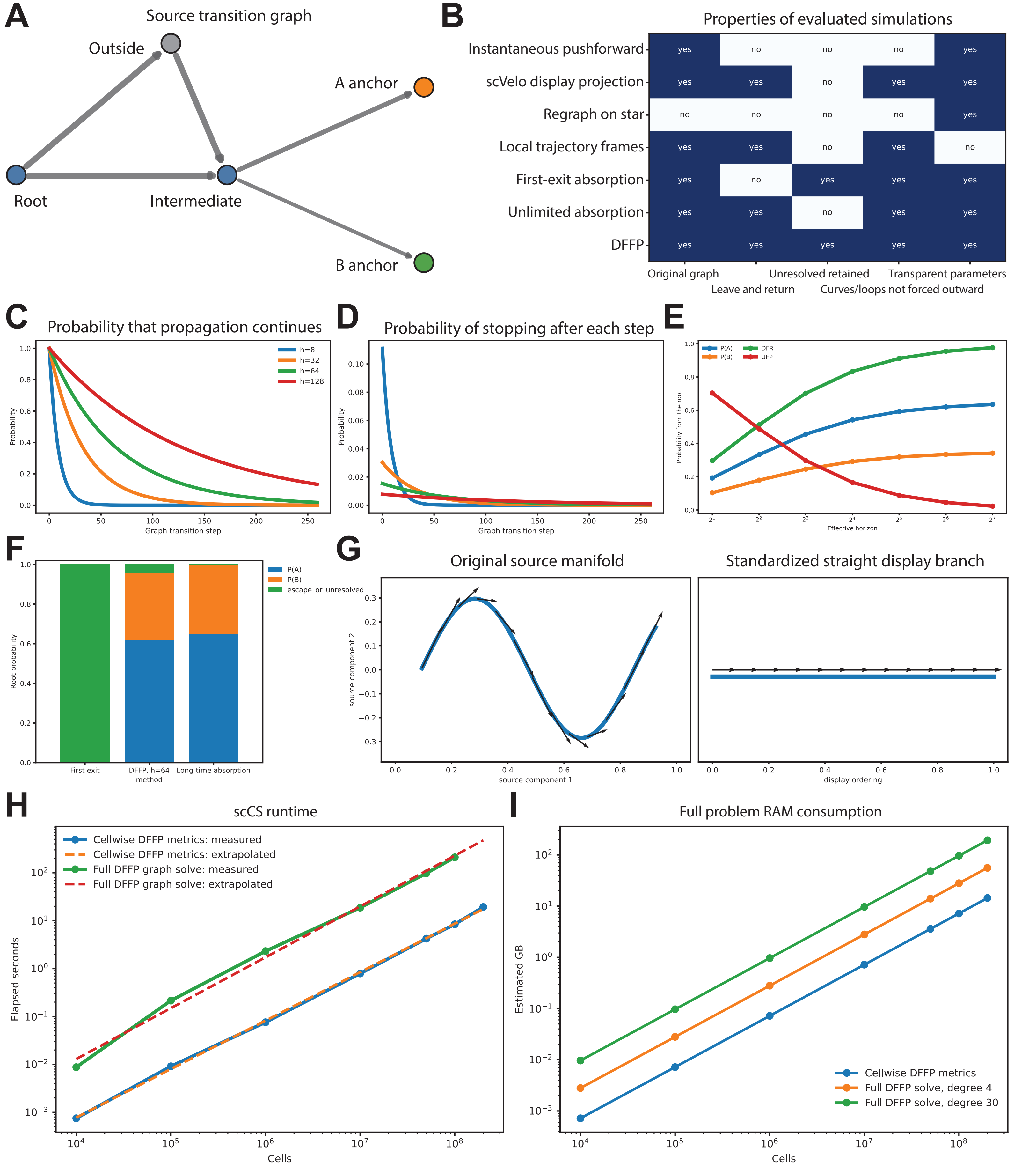
